# The interplay between supercoiling and DNA modifying enzymes at the

**DOI:** 10.1101/2025.10.02.680164

**Authors:** Elise M. Wilkinson, Antoine M. van Oijen, Timothy D. Craggs, Stefan H. Mueller, Lisanne M. Spenkelink

## Abstract

DNA supercoiling refers to the overwinding or underwinding of the double helix, forming plait-like structures called plectonemes. Supercoiling is involved in all biological processes that occur on DNA. Despite this involvement, single-molecule *in vitro* studies on these processes largely work with relaxed DNA, overlooking the role supercoiling is playing in these processes. We have constructed 18-kb linear DNA substrates that can be tethered to a microscope coverslip at multiple sites, allowing the template to be topologically constrained. We can then use an intercalating dye to induce positive or negative supercoiling, which we can observe at the single-molecule level using TIRF microscopy. We directly visualise plectonemes diffusing along the DNA. However, in the presence of the restriction enzyme EcoRV, plectonemes remain stationary at the EcoRV binding site. We show that EcoRV has an increased cleavage efficiency in the presence of supercoiling, even though its binding efficiency is unaffected. These findings reveal a relationship between DNA topology and the activity of DNA modifying enzymes. We hypothesise that supercoiling makes cleavage more energetically favourable for restriction enzymes that bend DNA. This work reveals the kinetic interplay between supercoiling and enzyme activity, highlighting how DNA itself can modulate the processes that act upon it.

**Graphical abstract:** 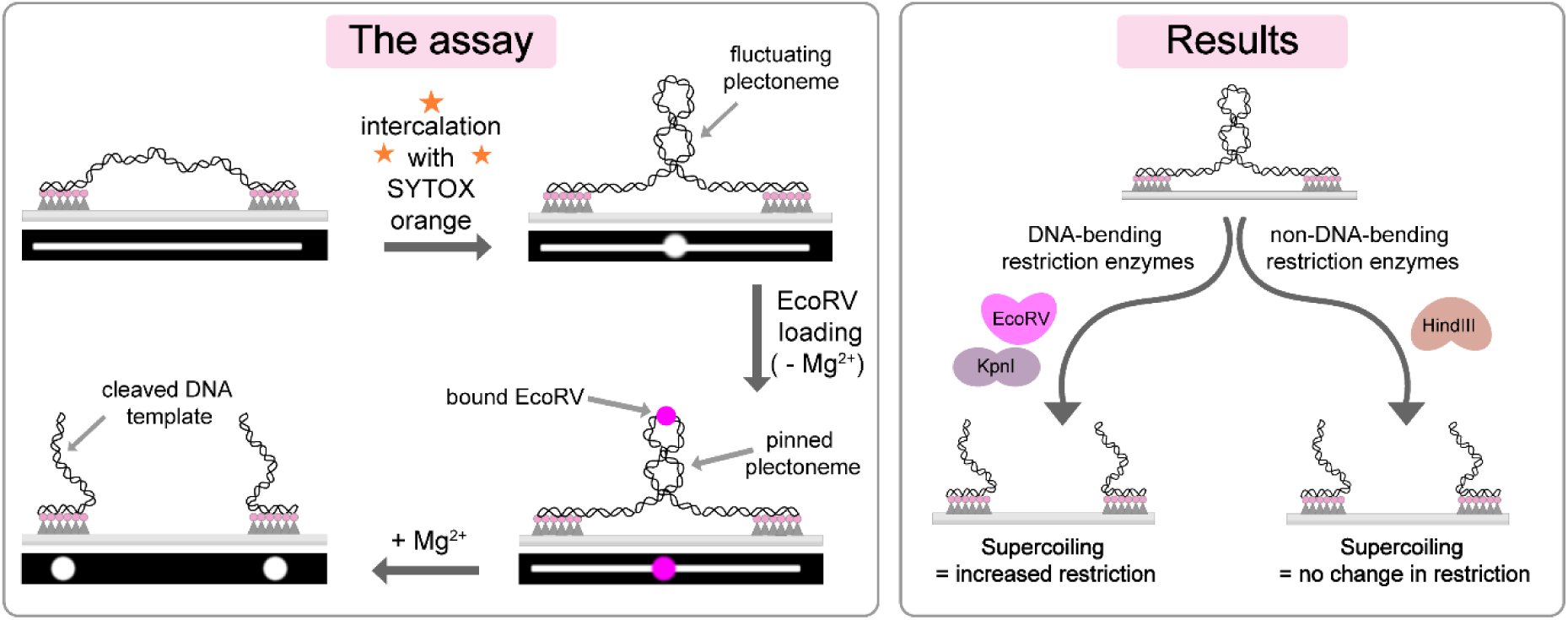

## Introduction

DNA supercoiling involves the over winding or under winding of double-stranded DNA, resulting in the formation of positively or negatively supercoiled structures. Supercoiling can exist in two forms, plectonemic or toroidal. Plectonemic supercoils, or plectonemes, are of particular interest for research as this form of supercoiling is present in cells across all domains of life and is therefore involved in all processes that act on DNA, notably, DNA replication and transcription. Despite the fact that supercoiling is involved in all processes that act on DNA *in vivo*, to date, single-molecule *in vitro* studies often work with relaxed DNA in the absence of supercoiling ^1–3^. Those that do consider supercoiling largely utilise magnetic and optical tweezer methods which do not allow for the direct observation of DNA kinetics and DNA-protein interactions in real time. Ultimately, this means that the role that supercoiling may be playing in these DNA processes is often overlooked and, when *in vitro* studies work with relaxed DNA, the physiological relevance of these studies may be limited.

Restriction enzymes present a simple model system to study the interaction of supercoiling with enzymes that act on DNA. This is due to restriction enzymes having distinct and easily quantifiable catalytic activity. The interactions of restriction enzymes with DNA can also be simply modulated through the selection of an enzyme with the desired number of cognate sites on the DNA substrate in use, as well as through the inclusion or exclusion of Mg^2+^, which is required as a cofactor for cleavage in type II restriction endonuclease reactions ^4,5^. We have chosen to use the type II restriction endonuclease, EcoRV, as a proof of concept due to EcoRV having well documented binding, bending, and cleavage activities ^6–11^. EcoRV is a member of the type IIP subclass of restriction endonucleases, which recognise palindromic sequences ^12^. EcoRV is derived from *E. coli* and, in nature, is responsible for protecting the bacterium from invading foreign DNA through cleavage at the centre of the GATATC sequence ^8,13,14^. In solution, EcoRV forms a homodimer. After dimer formation, EcoRV can then bind to the DNA at both cognate and non-cognate sites. In the case of the latter, the enzyme must then use facilitated diffusion to find the correct sequence. At the cognate site, EcoRV is able to induce a 44° DNA bend prior to cleavage ^7^. While the initial binding can be non-specific, EcoRV shows a very high specificity for bending and cleavage activities ^15–19^. While EcoRV is one of the most thoroughly studied restriction endonucleases, many studies work with relaxed DNA, and those that do use supercoiled DNA often do not consider supercoiling or its role in enzyme behaviour. Furthermore, many of the studies that do work with supercoiled DNA use ensemble-averaging methods, which are insensitive to transient interactions and conformational dynamics of DNA. While these studies have provided valuable information, the results are static in nature, with kinetics having to be inferred, rather than observed ^20–23^. As supercoiling is such a dynamic process, we believe that directly observing these kinetics is vital to truly understanding how supercoiling influences, or modulates, the activity of enzymatic processes. Studying supercoiling using single-molecule methods aids us in disentangling kinetic details and enzyme behaviours, allowing for the separation of enzyme binding from endonuclease activities, as well as allowing us to watch, in real-time, the co-localisation of supercoiled structures with bound enzymes.

Recently, a single-molecule supercoiling assay has been developed, allowing for the visualisation of plectonemes in the presence and absence of plectoneme-pinning agents ^24–28^. This single-molecule supercoiling assay allows for the visualisation of supercoiling dynamics, both in the absence and presence of DNA-modulating enzymes, such as EcoRV, using total internal reflection fluorescence (TIRF) microscopy. Additionally, this assay allows us to observe how enzyme function is affected by different supercoiled states, helping us to further understand the question – is supercoiling merely a by-product of the processes that act on DNA, or does supercoiling itself act as a regulatory mechanism for these processes?

## Materials and methods

### Torsionally constrained template construction

Template construction protocols were adapted from ^29^. Briefly, an 18.3-kb plasmid (pUBER) purified by Aldevron (USA) was digested with BstXI (1 unit/µg DNA) for 10 hours at 37 °C in 1x buffer 3.1 (New England Biolabs), followed by a heat inactivation at 80 °C for 20 minutes (see Supplementary Figure S1 online).

Capping oligonucleotides were designed and purchased from Integrated DNA Technologies (IDT) (see Supplementary Table S1 online). Oligonucleotides were annealed at a 1:1 molar ratio to generate two different 6-biotin-containing blocks, one with a Cy5 fluorophore for directionality, and one without. These oligo blocks were ligated onto either end of the 18.3-kb digested pUBER fragment with T4 DNA ligase (100 units/µg DNA) in 1x Cutsmart buffer supplemented with 1 mM ATP.

Excess oligonucleotide and T4 DNA ligase were removed by loading onto a Sepharose 4B column (1 cm x 25 cm) in the presence of 12 mM EDTA and 300 mM NaCl. FPLC fractions were measured on a nanodrop, and fractions with a DNA concentration greater than 10 ng/µL were kept for use in single-molecule experiments.

### Flow cell construction and microscope set up

Microfluidic flow cells were prepared as outlined previously ^30,31^. Briefly, a polydimethylsiloxane (PDMS) flow chamber containing three parallel channels was placed on top of a PEG-biotin-functionalised microscope coverslip (see Supplementary Figure S2 online). Polyethylene tubing (BTPE-60, Instech Laboratories) was inserted into the entrance and exit of the PDMS channels. A syringe pump (Adelab Scientific, Australia) was connected to the outlet tubing to allow for buffer flow. To help prevent non-specific interactions of proteins and DNA with the surface, the chamber was blocked with buffer containing 20 mM Tris-HCl pH 7.5, 2 mM EDTA, 50 mM NaCl, 0.2 mg/ml BSA, and 0.005% Tween-20. The flow cell was placed on an inverted microscope (Nikon Eclipse Ti-E) with a CFI Apo TIRF 100x oil-immersion TIRF objective (NA 1.49, Nikon, Japan). All experiments were carried out at room temperature.

Double-stranded DNA was visualised in real-time through SYTOX orange staining (Invitrogen) which was excited by a 514-nm laser (Coherent, Sapphire 514-150 CW) at 1.35 W cm^−2^. A 647-nm laser (Coherent, Obis 647–100 CW) at 12.5 W cm^−2^ was used to image Cy5 DNA end labels and fluorescently labelled EcoRV (BG-AF647).

### Intercalation-induced supercoiling

Supercoiling was induced and visualised using the intercalating dye, SYTOX Orange (Thermo Fisher Scientific), which transiently binds between the base pairs of double-stranded DNA and induces twist upon topologically constrained DNA templates. Positive and negative supercoiling was selected for by increasing or decreasing the SYTOX orange concentration from DNA loading to DNA imaging. That is, positive supercoiling was induced when the DNA was loaded into the flow cell and bound to the coverslip in the presence of a lower SYTOX orange concentration, then imaged in the presence of a higher SYTOX orange concentration. The opposite is true for negative supercoiling. As supercoiling is induced by changing the SYTOX orange concentration between DNA loading and DNA imaging, a ‘constrained control’ was achieved by maintaining a constant SYTOX orange concentration between DNA loading and DNA imaging. In this case, no supercoiling was induced, which allowed for the same DNA template to be used as a non-supercoiled control.

### Modulating degree of supercoiling

In addition to modulating positive and negative supercoiling via SYTOX orange concentration, the degree of supercoiling can also be modulated in this assay. Two degrees of supercoiling were established – ‘high supercoiling’ and ‘low supercoiling’. These degrees of supercoiling were selected for by modulating the flow rate during DNA loading which, in turn, modulated the degree of DNA stretch and the amount of ‘slack’ DNA available for plectonemes to form (see Supplementary Figure S3 online). In this study, DNA loading flow rates of 10 µL/min and 20 µL/min were used for ‘high’ and ‘low’ degrees of supercoiling respectively.

### Positive supercoiling assay

DNA templates were loaded into the flow cell in 500 µL of magnesium-free supercoiling imaging buffer (80 mM Tris pH 7.5, 100 mM NaCl, 5 mM CaCl_2_) containing no SYTOX orange at a flow rate of 10 µL/min for high supercoiling conditions, and 20 µL/min for low supercoiling conditions (see Supplementary Figure S3 online). An additional 200 µL of imaging buffer containing 250 nM SYTOX orange and supplemented with photo-protective reagents (4mM Trolox, 2.25 mM protocatechuic acid (PCA), and 50 nM nuclease-free protocatechuate-3,4-dioxygenase (scientific reports) (Oriental Yeast Co., Ltd)) was then loaded at 10 µL/min to induce positive supercoiling. Data were collected through continuous imaging for 3 minutes in the absence of flow with an exposure time of 200 ms. Four 82 µm x 82 µm fields of view were imaged for each experiment.

### Negative supercoiling assay

The above was followed, except DNA templates were loaded into the flow cell in the presence of 500 nM SYTOX orange. Negative supercoiling was then induced by the loading of an additional 200 µL of imaging buffer containing 250 nM SYTOX orange and supplemented with photo-protective reagents (4 mM Trolox, 2.25 mM PCA, and 50 nM nuclease-free PCD). Data were collected through continuous imaging for 3 minutes in the absence of flow with an exposure time of 200 ms. Four 82 µm x 82 µm fields of view were imaged for each experiment.

### Constrained control assay

The above was followed, except DNA templates were loaded into the flow cell in the presence of 250 nM SYTOX orange. DNA loading flow rates of 10 µL/min and 20 µL/min were used, depending on the degree of ssupercoiling desired. For imaging, an additional 200 µL of imaging buffer containing 250 nM SYTOX orange and supplemented with photo-protective reagents (4 mM Trolox, 2.25 mM PCA, and 50 nM nuclease-free PCD) was loaded. Since there was no change in SYTOX orange concentration, no supercoiling was induced. Data were collected through continuous imaging for 3 minutes in the absence of flow with an exposure time of 200 ms. Four 82 µm x 82 µm fields of view were imaged for each experiment.

### Fluorescent labelling of EcoRV

We designed an EcoRV-SNAP-tag-Flag-tag fusion construct for easy labelling and purification which can be rapidly expressed using an *in vitro* transcription translation (IVTT) reaction following company instructions (New England Biolabs) ^32^ (see Supplementary Table S2 online). The construct was synthesized as a gene block (Integrated DNA Technologies). Following expression through IVTT, 2 µL of SNAP-Surface Alexa Fluor 647-nm dye (New England Biolabs) was added to the sample and incubated at 37 °C for 30 min. The reaction product was then purified using anti-flag magnetic beads (Sigma-Aldrich). Enzyme purity was confirmed by SDS-PAGE and activity tests confirmed that the fluorescently labelled EcoRV was comparable in activity to commercial EcoRV (New England Biolabs) (see Supplementary Figure S4 online). The fluorescently labelled EcoRV was and then diluted in EcoRV storage buffer as outlined by New England Biolabs (10 mM Tris-HCl, 200 mM NaCl, 1 mM DTT, 0.1 mM EDTA, 200 µg/ml BSA, 50% Glycerol, pH 7.4 @ 25°C) and aliquoted and stored at -20°C.

### Fluorescently labelled EcoRV photobleaching assay

To determine the fluorescence intensity of a single EcoRV dimer, 647-EcoRV was diluted 1000x and approximately 40 µL was sandwiched between two microscope coverslips. This coverslip was placed on the microscope and two fields of view were selected. These fields of view were illuminated by a 647-nm laser at 12.5 W cm^−2^ and imaged continuously with an exposure of 200 ms until complete photo-bleaching was observed. These data were then processed and analysed using an in-house script. From this, we obtained the average intensity of two 647-fluorophores, corresponding to one EcoRV dimer (see Supplementary Figure S4 online).

### Plectoneme pinning assay with fluorescently labelled EcoRV

The microfluidic flow cell was prepared, and the supercoiling assay was set up as outlined above, depending on if positive or negative supercoiling was desired. The presence of supercoiling was confirmed through brief excitation with a 514-nm laser (Coherent, Sapphire 514-150 CW) at 400 mW cm^−2^. Following the confirmation of supercoiling, two fields of view were selected and saved. In these two fields of view, the Cy5 end label of the DNA template was imaged to determine the orientation of DNA templates. End labels were then photo-bleached by continuous excitation with a 647-nm laser to prevent signal overlap with the labelled EcoRV. Following this, fluorescently labelled EcoRV was loaded into the flow cell in 200 µL of imaging buffer (80 mM Tris pH 7.5, 100 mM NaCl, 5 mM CaCl_2_) supplemented with 250 nM SYTOX orange and photo-protective reagents (4 mM Trolox, 2.25 mM PCA, and 50 nM nuclease-free PCD) at a flow rate of 10 µL/min. After EcoRV loading, the same fields of view were selected, and double-stranded DNA and 647-EcoRV were illuminated by 514-nm and 647-nm lasers concurrently and imaged in the absence of flow for six minutes at 0.5 frames per second with an exposure time of 200 ms. The fluorescence intensity of a single EcoRV dimer was used to determine the number of EcoRV dimers bound to a DNA molecule. These values were used to determine the binding efficiency of EcoRV in different states of supercoiling.

### EcoRV cleavage assay

DNA templates were loaded as described above, depending on the supercoiled state required. Following the loading of SYTOX orange in magnesium-free supercoiling imaging buffer, fluorescently labelled EcoRV was loaded in an additional 200 µL of imaging buffer supplemented with photo-protective agents (4 mM Trolox, 2.25 mM PCA, and 50 nM nuclease-free PCD) and 250 nM SYTOX orange at a flow rate of 10 µL/min. Four fields of view were selected and saved. Cy5 end labels were excited by a 647-nm laser (Coherent, Obis 647–100 CW and imaged to obtain directional information. The double stranded DNA was excited by a 514-nm laser (Coherent, Sapphire 514-150 CW) and imaged briefly. These images served as a ‘before cleavage’ DNA count. 200 µL of magnesium-containing cleavage buffer (80 mM Tris pH 7.5, 100 mM NaCl, 5 mM MgCl_2_) supplemented with photo-protective agents (4 mM Trolox, 2.25 mM PCA, and 50 nM nuclease-free PCD) and 250 nM SYTOX orange was then loaded at a flow rate of 10 µL/min. After flow had stopped, the original four fields of view were again selected, and the double-stranded DNA remaining was excited by the 514-nm laser and imaged briefly. These images served as an ‘after cleavage’ DNA count.

### KpnI and HindIII cleavage assays

The above protocol for the EcoRV cleavage assay was followed, with KpnI-HF and HindIII-HF (0.1 U/µL cleavage buffer) (New England Biolabs) substituted as required. As KpnI and HindIII were not fluorescently labelled, plectoneme pinning and binding efficiency were not considered.

It should be noted that all cleavage assays were carried out in the magnesium-containing cleavage buffer (80 mM Tris pH 7.5, 100 mM NaCl, 5 mM MgCl₂), rather than in the optimal reaction buffers recommended by New England Biolabs. Therefore, we focused on comparing the relative cleavage efficiency of each restriction enzyme between different supercoiled states and did not consider the overall lower activity across all states to be a concern.

### Cross-flow experiments

The microfluidic flow cell was constructed as outlined above however, instead of three parallel channels, the PDMS contained two intersecting channels forming a cross (see Supplementary Figure S2 online). These channels were blocked and washed as described above. Following this, DNA templates were loaded into one channel (flow direction one) in a total of 500 µL magnesium-free supercoiling buffer at a flow rate of 10 µL/min. During DNA loading, the tubing of the second channel was clamped to prevent unwanted sideways flow that may occur due to the intersecting channels. Following DNA loading, 200 µL of imaging buffer supplemented with 250 nM SYTOX orange and photo-protective agents (4 mM Trolox, 2.25 mM PCA, and 50 nM nuclease-free PCD) were loaded into this same channel at a flow rate of 10 µL/min. Supercoiling was confirmed through brief illumination with a 514-nm laser. Cy5 end labels were imaged and photo-bleached through continuous illumination with a 647-nm laser. If 647-EcoRV was used, it was then loaded into the first channel as outlined above for regular 647-EcoRV experiments.

The first channel was then clamped and disconnected from the syringe pump, and the second channel was connected. Imaging buffer supplemented with 250 nM SYTOX orange was then loaded into this second channel (flow direction two) at flow rate of 60 µL. Double-stranded DNA and 647-EcoRV were illuminated by 514-nm and 647-nm lasers concurrently and imaged under buffer flow for six minutes at 0.5 frames per second with an exposure time of 200 ms.

### SYTOX orange intensity during overwinding and underwinding

Supercoiling experiments in the absence of DNA-binding enzymes were carried out as outlined above. Following data collection, movies of supercoiled molecules were categorised depending on whether they relaxed due to photo-nicking or remained supercoiled within the three minutes of imaging.

In total, 22 negatively supercoiled molecules that relaxed due to photo-induced nicking were collected. These molecules were then analysed in ImageJ to measure the integrated SYTOX orange intensity of every frame of each molecule’s movie. The frame in which relaxation occurred was noted for all molecules, and intensities were compiled into plots, where t=0 corresponded to the photo-nicking event. Average SYTOX orange intensity before and after nicking was then determined and plotted.

### Data analysis

All background corrections, drift corrections, and analyses were carried out using ImageJ/Fiji (1.51w) and in-house plugins, found here: https://github.com/Single-molecule-Biophysics-UOW.

### Statistical analysis

Statistical analysis of data was carried out using ImageJ/Fiji (1.51w), Microsoft Excel 2016, and Spyder (Anaconda 3) using in-house scripts. Data were collected from images from at least two independent experiments for each condition. Average z-projections of each field of view were used for calculation of cleavage and binding efficiencies. N values correspond to the number of field of views analysed.

## Results

### Visualising positive and negative supercoiling at the single-molecule level

To visualise supercoiling at the single-molecule level, long linear DNA templates need to be immobilised to a microscope coverslip in a way that allows induction of supercoiling. Protocols for the immobilisation of long linear DNA templates on to the surface of a microscope coverslip through biotin-streptavidin linkages are well established ^29,33–39^. However, these DNA templates contain one biotin on each end, usually attached via linker moieties which allow rotation around the radial axis. This does not allow for the induction or visualisation of supercoiling as any rotational stress on the DNA would be alleviated through rotation around the DNA axis.

Recently, this has been overcome by the development of a single-molecule supercoiling assay that uses multiple biotins at each end of a long linear DNA template. These multiple biotins allow for the DNA template to be tethered and constrained to a functionalised coverslip, allowing for supercoiling to be induced upon it ^24–28^. Similarly, we created 18-kb long linear DNA templates that contained six internal biotins at each end (Figure 1A).

**Figure 1:**
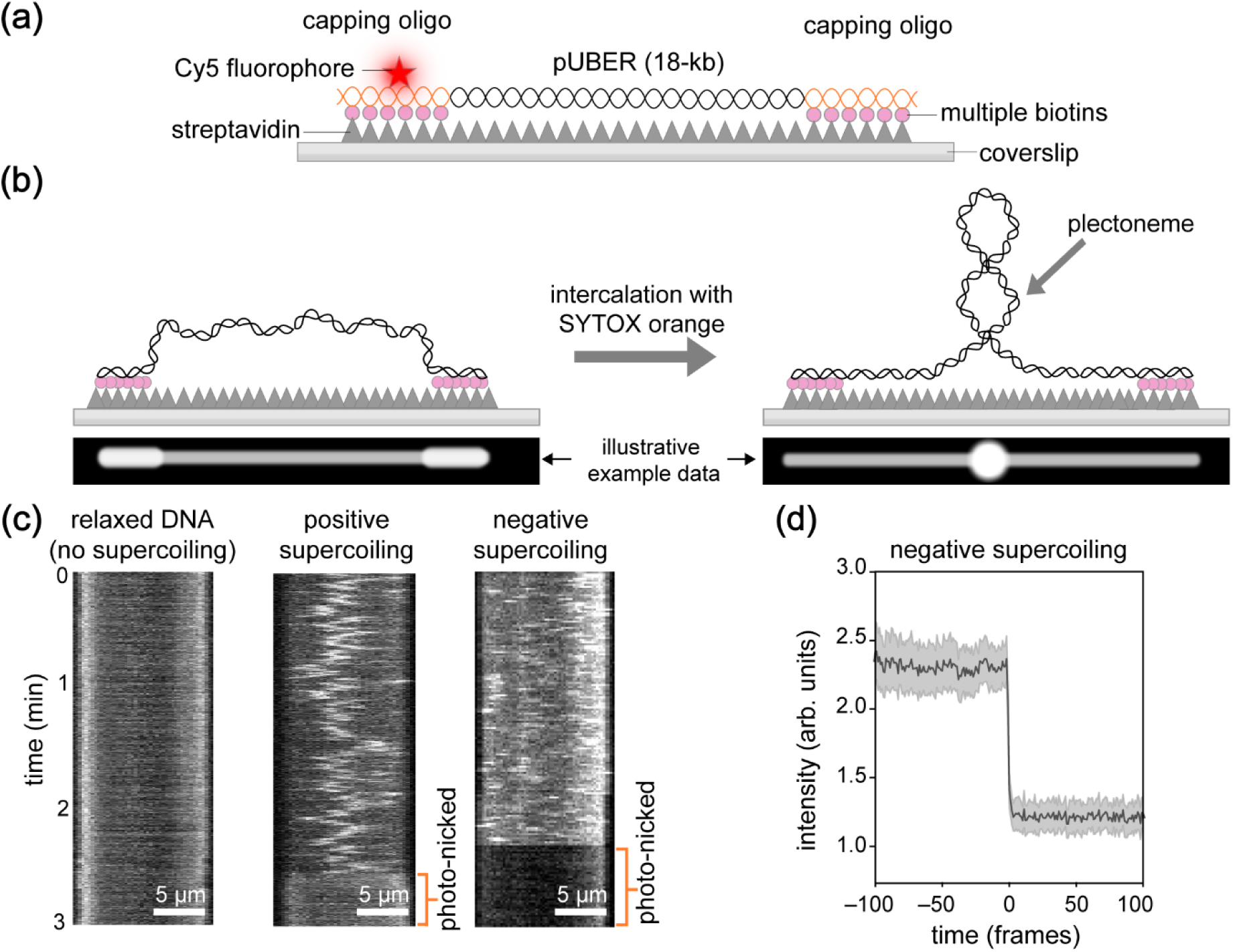
single-molecule supercoiling assay using TIRF microscopy. **(a)** Schematic depicting the 18-kb linear DNA template containing six internal biotins on each end, which allow the template to be topologically constrained to a coverslip. **(b)** Schematics depicting the bound and constrained DNA template prior to (left) and following (right) intercalation with SYTOX orange. The illustrative example data represents what this would look like when imaged using TIRF microscopy. **(c)** Three kymographs depicting the DNA template in the absence of supercoiling (left), in the presence of positive supercoiling (middle), and in the presence of negative supercoiling (right). Kymographs consist of every frame of a 3-minute movie (200 ms exposure) condensed into a single image, with time going from top to bottom. Scale bars = 5 µm. **(d)** Graph depicting SYTOX orange intensity before and after negatively supercoiled DNA molecules relax due to laser-induced photo-nicking.

DNA templates were loaded into a microfluidic flow cell containing a streptavidin-functionalised coverslip. Through the laminar flow, DNA templates become doubly tethered to the coverslip though multiple biotin-streptavidin linkages. Flow rates were adjusted during DNA loading to modulate the degree of stretch, and hence the degree of supercoiling (see methods).

To induce and visualise supercoiling, we used the intercalating dye SYTOX orange (Thermo Fisher Scientific), which transiently binds between base pairs, inducing twist and hence, supercoiling (Figure 1B) ^24^. Plectonemes appeared as bright local fluorescent spots that fluctuated along the DNA template and disappeared upon photo-induced nicking (Figure 1C). The appearance and behaviour of these fluctuating plectonemes was consistent with previous studies ^24–28^.

During the acquisition, templates appear to spontaneously relax (Figure 1C). This relaxation is due to photo nicking — a laser-induced form of DNA damage that introduces single-stranded nicks in the DNA ^40^. When measuring the fluorescence intensity of negatively supercoiled DNA molecules before and after photo nicking, we observed a decrease in fluorescence intensity after photo nicking. This indicates that the under wound negatively supercoiled DNA allows for increased SYTOX orange binding whilst the molecule is in its supercoiled state (Figure 1D). These results are consistent with previous findings ^25^ and confirm that, in our assays, SYTOX orange intercalation is in fact inducing a topological change that can then, in turn, influence further intercalation of SYTOX orange. Notably, this altered binding activity is then reverted to that of a non-topologically constrained molecule upon relaxation due to photo-damage (Figure 1D).

### Plectonemes are pinned to occupied EcoRV restriction sites

Plectoneme pinning can occur because of a number of reasons, including, but not limited to, sequence preference, DNA bends, bound proteins, and DNA lesions. Our single-molecule supercoiling assay has the potential to accommodate for a number of these plectoneme pinning conditions, however we chose to start with the type II restriction endonuclease, EcoRV, as an easy model system for plectoneme pinning. EcoRV was selected because of its specific binding and cleavage activity, as well as its well-established DNA bending activity ^6–10,15^. As EcoRV requires magnesium to cleave, we can separate EcoRV binding activity from its cleavage activity through the exclusion or inclusion of MgCl_2_ (Figure 2A). To focus on its binding activity first, we excluded MgCl_2_ and instead supplemented the buffer with CaCl_2_.

**Figure 2:**
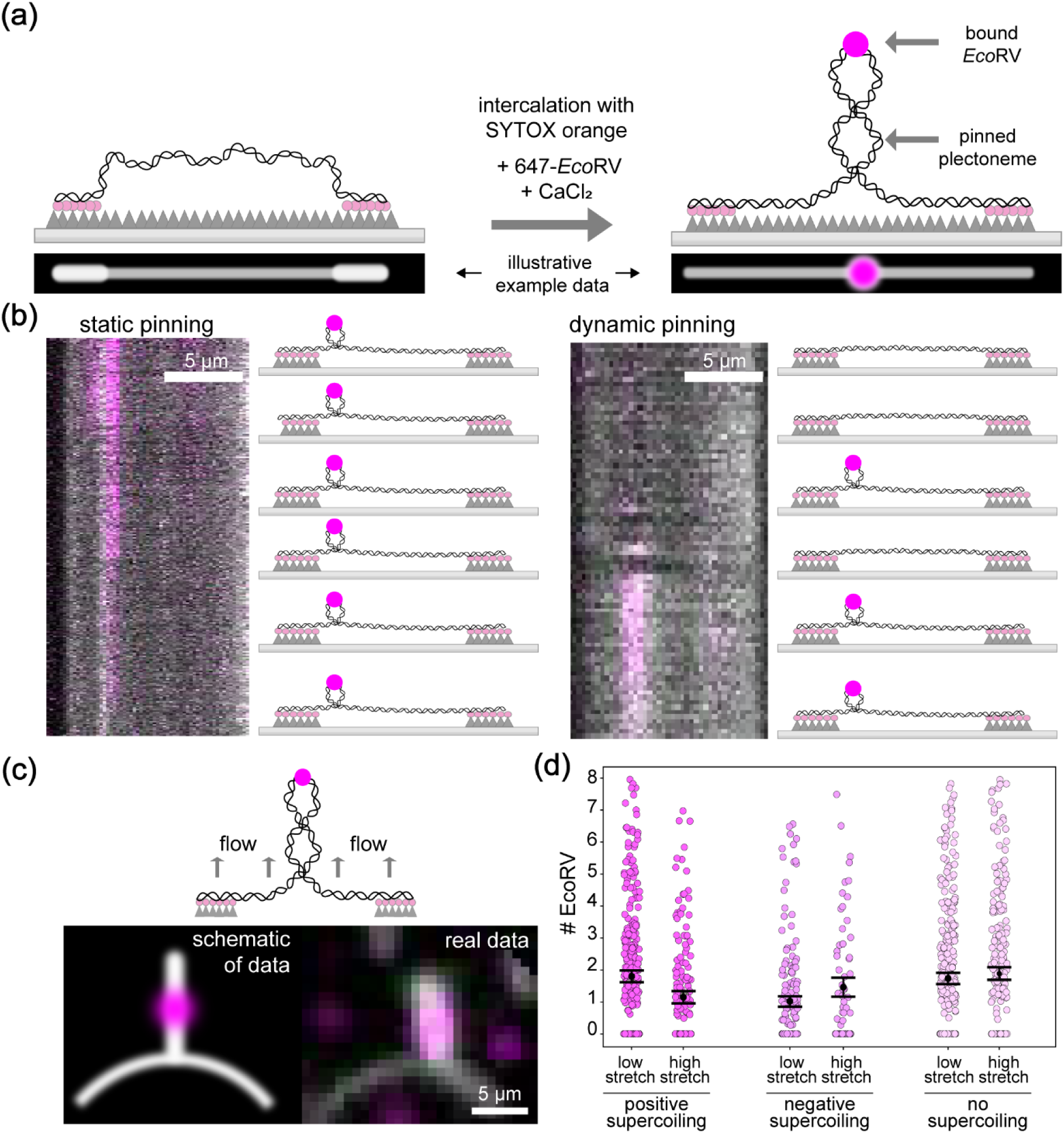
EcoRV-induced plectoneme pinning at the single-molecule level. **(a)** Schematics depict the DNA template prior to intercalation with SYTOX orange (left) and following intercalation and EcoRV-induced pinning (right). The illustrative example data represents how this would appear when imaged using TIRF microscopy. **(b)** Kymographs of positively supercoiled molecules showing two different plectoneme pinning event types, with corresponding schematics representing the binding and plectoneme-inducing events that are occurring. Kymographs consist of every frame of a 6-minute movie (200 ms exposure, 500 ms delay) condensed into a single image with time going from top to bottom. Scale bars = 5 µm. **(c)** Schematic representing the supercoiling cross-flow assay in the presence of 647-EcoRV, in addition to both illustrative and real data showing EcoRV colocalised with a pinned plectoneme. **(d)** Scatter plot presenting the number of EcoRV sites occupied per DNA molecule under different degrees of supercoiling. Error bars represent standard error of the mean (SEM).

We expressed snap-tagged EcoRV in a cell-free expression system (in vitro transcription-translation) ^41,42^ and labelled it with a fluorescent dye. This labelled EcoRV was then included in the established supercoiling assay to look for co-localisation between plectonemes and bound EcoRV dimers.

After loading EcoRV we observed stationary bright fluorescent spots, or plectonemes, colocalised with labelled-EcoRV signal (Figure 2A). A number of event types were observed, of which, two were prominent.

Figure 2B (left) shows static plectonemes colocalised with stably bound EcoRV for the duration of imaging. The molecule in the kymograph on the right shows no plectonemes until EcoRV binding occurs, where we then see a plectoneme form at this binding site. EcoRV binds on and off, with the plectoneme having corresponding intensity. While we do see a range of different event types, overall, our results are consistent with a model that suggests that EcoRV-binding induces plectoneme formation.

Additionally, we can look at these pinned plectonemes under cross flow, where we can clearly see co-localisation between the plectoneme and the bound 647-EcoRV (Figure 2C). From this, we can also confirm that the bound EcoRV is positioned within the plectoneme.

Next, we aimed to quantify the number of EcoRV bound to each DNA template, under different levels of supercoiling. We can induce different levels of supercoiling by modulating the flow rate during DNA loading (see methods). Because each EcoRV monomer is labelled with one fluorophore, we can use the fluorescence intensity to count the number of EcoRV dimers bound to each DNA molecule (see Supplementary Figure S4 online). Each DNA molecule has a maximum of eight sites available (see Supplementary Figure S1 online). We for all supercoiling conditions we found approximately two EcoRV dimers per DNA molecule (Figure 2D).

For comparison, we also measured the number of EcoRV molecules bound to constrained, but not supercoiled DNA molecules. To set up these measurements, we used the same flow rates during DNA loading, to ensure that the level of stretch was the same as in our supercoiling conditions. However, we did not change the SYTOX orange concentrations from DNA loading to imaging, ensuring that supercoiling is not induced. Again, we found ∼2 EcoRV dimers per DNA molecule. These results show that EcoRV binding is not affected by supercoiling.

### Positive and negative supercoiling modulates the cleavage activity of EcoRV

The results described thus far have been obtained in the absence of magnesium, allowing us to observe the binding activity of EcoRV, whilst preventing cleavage from occurring. To look at the cleavage activity of EcoRV, we now include MgCl_2_ (Figure 3A).

**Figure 3:**
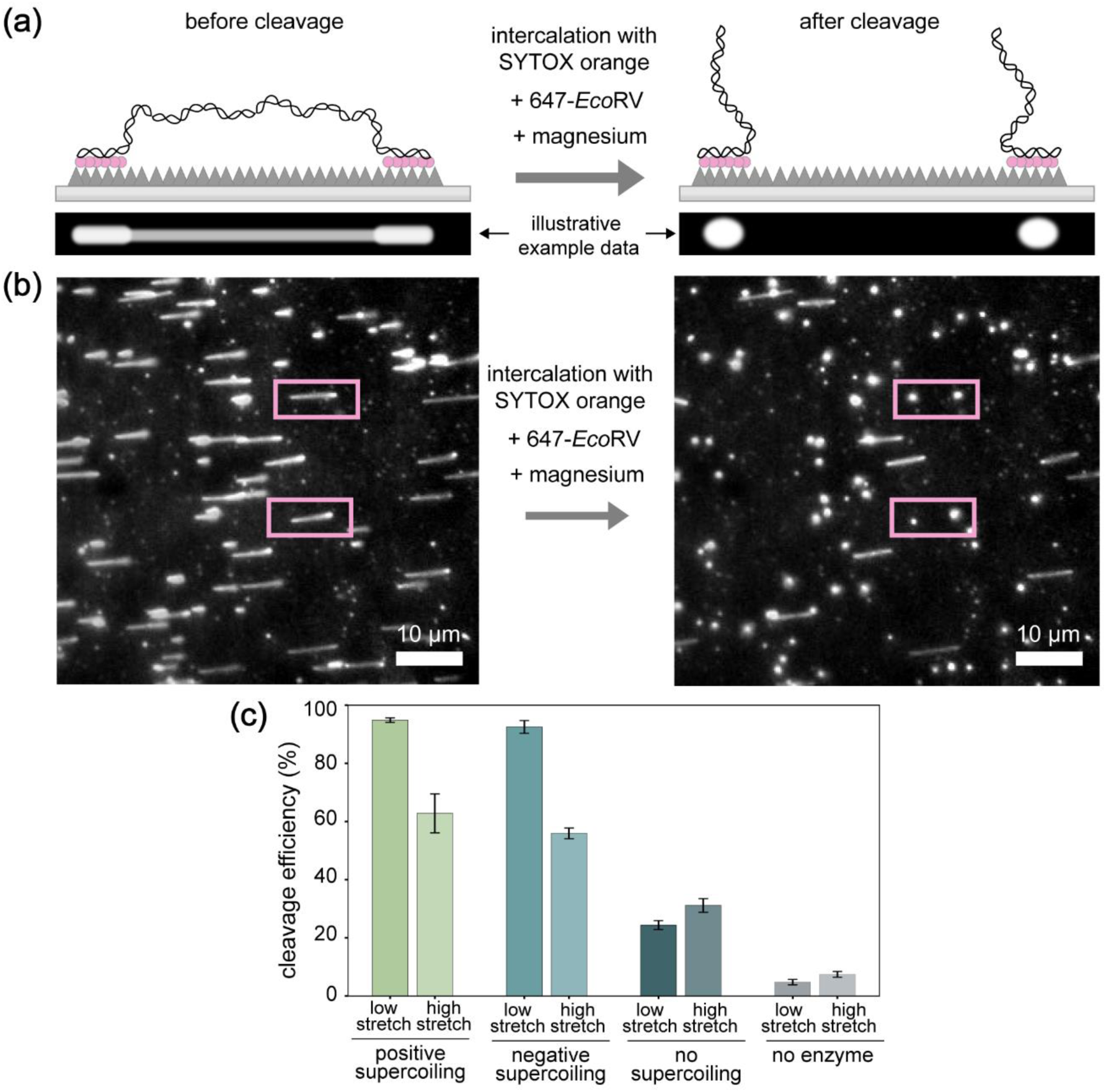
EcoRV cleavage assay in the presence of different degrees of positive and negative supercoiling. **(a)** Schematic and illustrative data representing the constrained DNA template before and after cleavage by EcoRV, which is induced through the introduction of Mg^2+^. **(b)** Representative fields of view showing a field of view before and after cleavage by EcoRV. Molecules were highly positively supercoiled, the condition in which we observed the highest cleavage efficiency by EcoRV. Scale bars = 10 µm. **(c)** Bar graph presenting EcoRV cleavage efficiency in the presence of different degrees of supercoiling, which was modulated through DNA loading flow rate and SYTOX orange concentration. Errorbars represent standard deviation (σ).

Figure 3B shows a representative field of view before induction of supercoiling (left) and after induction of a high degree of positive supercoiling and introduction of EcoRV in the presence of MgCl_2_ (right). Under these conditions of positive supercoiling, we find that 94.9 ± 1.9 % (n = 12) of DNA templates are cleaved. In the presence of a lower degree of positive supercoiling, this cleavage efficiency dropped to 62.8 ± 11.6% (n =12) (Figure 3C). Similarly, in the presence of a high degree of negative supercoiling, it was found that EcoRV had a cleavage efficiency of 92.5% ± 4.4% (n =12). In the presence of a lower degree of negative supercoiling, this cleavage efficiency dropped to 55.9 ± 4.9% (n = 12).

Surprisingly, in the absence of supercoiling, the cleavage efficiency dropped almost by a factor of 4 to 24.4 ± 4% (n = 12).To test if these differences in cleavage efficiency are a result of supercoiling, and not simply a result of stretch, we looked at the cleavage activity of EcoRV in the absence of supercoiling at high and low-stretch conditions, which did not yield significant differences (high-stretch: 24.4 ± 4% (n = 12), low stretch: 31.1 ± 6.7% (n =12)). (Figure 3C).

Additionally, we looked at the degree of DNA template cleavage and/or breakage in the absence of any restriction enzymes in order to ensure that the cleavage we observe is not a result of flow or laser induced breakage. In the presence of a high degree of positive supercoiling, we observed that 4.8 ± 2.1% (n = 8) of molecules cleaved/broke. Similarly, in the presence of a low degree of positive supercoiling, we observed that 7.4 ± 2.2% (n = 8) of molecules cleaved/broke (Figure 3C). These findings indicate that the cleavage activity of EcoRV is promoted by both positive and negative supercoiling, and that the degree of DNA stretch has no effect.

From these results, we can conclude that we have observed, for the first time at the single-molecule level, that both positive and negative supercoiling are modulating the cleavage activity of EcoRV. These results are consistent with previous ensemble studies ^20^, where it was found that supercoiling assisted EcoRV in finding its cognate sequence through 3D diffusion. However, this study does not consider the significant bending activity of EcoRV and whether this may play a role in its modulation by supercoiling.

### Positive and negative supercoiling regulates the cleavage activity of DNA-bending restriction enzymes

From our results, we determined that the cleavage activity of EcoRV is modulated by both positive and negative supercoiling. We hypothesise that this modulation in cleavage efficiency is linked to the bending of DNA. That is, in the presence of supercoiling, it is more energetically favourable for DNA-bending restriction enzymes to bend and cleave DNA. So, to see if this holds true for other type II restriction enzymes, we have selected two additional enzymes, one that only bends DNA minimally, and one that induces a significant bend, similar to EcoRV. As EcoRV induces a 44° bend at its DNA binding site ^7^, we decided to select HindIII, which only bends DNA minimally ^43^, and KpnI, which induces a ∼35° bend ^44^. It should be noted that these three enzymes do not have the same number of cognate sequences within our DNA template (see Supplementary Figure S1 online). Therefore, direct comparisons between enzymes are not meaningful. Instead, we compare the activity of each enzyme under different supercoiling conditions relative to its own activity on relaxed DNA. The activity of these restriction enzymes were confirmed through a gel-based activity test prior to commencing single-molecule assays (see Supplementary Figure S5 online).

Figure 4A and 4B show representative fields of view showing the number of linear molecules before and after addition of enzyme in the presence and absence of supercoiling. We can see that in the presence of a high degree of positive supercoiling, HindIII had a cleavage efficiency of just 11.8 ± 5.3% (n= 13) (Figure 4A,C). Similarly, in the absence of supercoiling, this cleavage efficiency was 12.2 ± 5 % (n = 13). These results indicate that HindIII cleavage efficiency is not influenced by supercoiling.

Contrastingly, we see that in the presence of a high degree of positive supercoiling, KpnI has a cleavage efficiency of 46.5 ± 5.1% (n = 12) (Figure 4B,C). In the absence of supercoiling, this cleavage efficiency then drops to 14.8 ± 2.2% (n = 12). These results align with the results obtained for EcoRV, where we see that the cleavage efficiency is positively modulated by the presence of supercoiling (Figure 4C).

**Figure 4.**
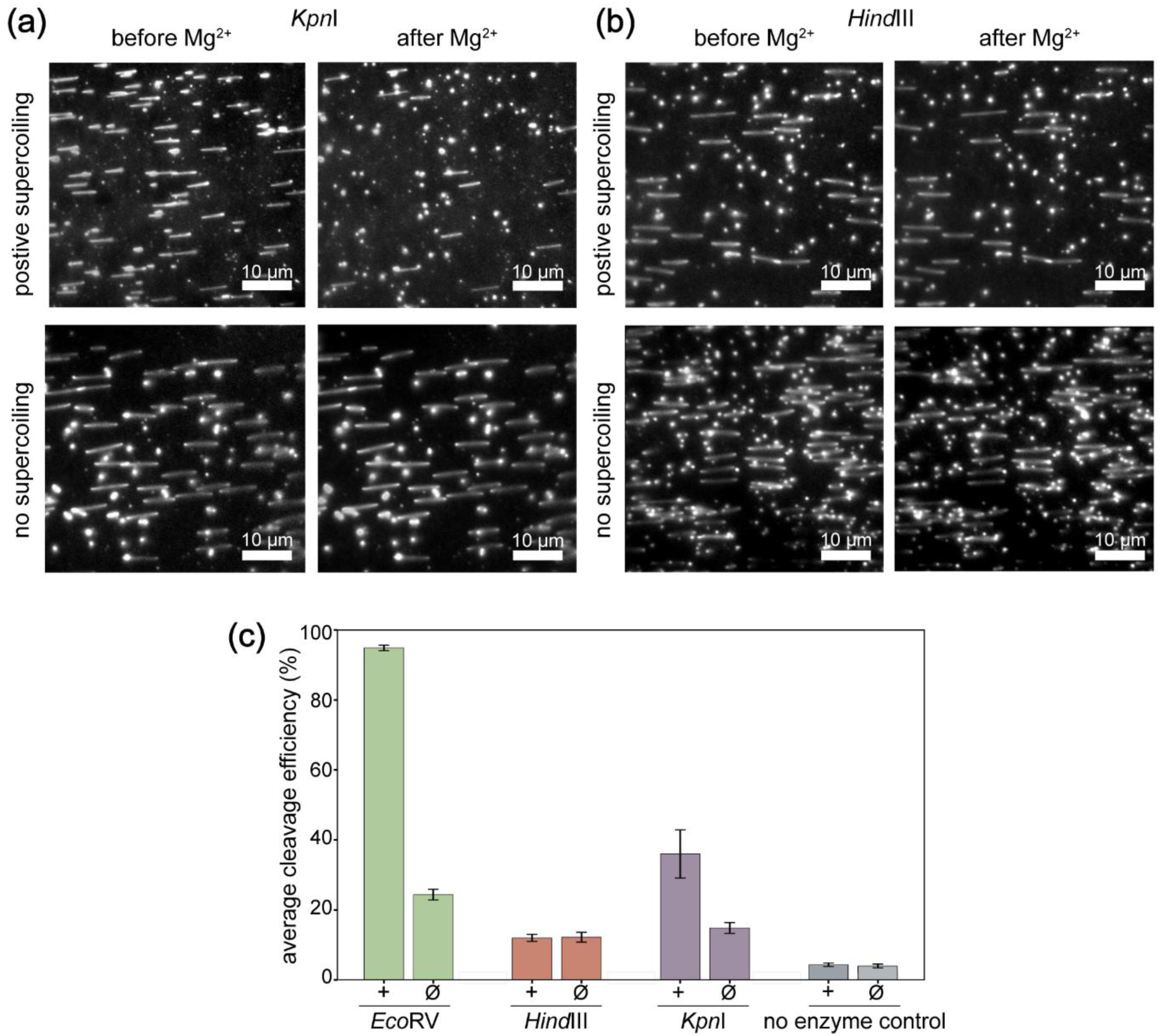
The cleavage efficiency of the DNA-bending enzyme, KpnI, and non-DNA-bending enzyme, HindIII, in the presence of different degrees of supercoiling. **(a)** Example field of views showing the cleavage activity of HindIII in the presence of positive supercoiling compared to the cleavage efficiency in the absence of supercoiling. Scale bars = 10 µm. **(b)** Example field of views showing the cleavage activity of KpnI in the presence of positive supercoiling compared to the cleavage efficiency in the absence of supercoiling. Scale bars = 10 µm. **(c)** Bar graph depicting the cleavage effiencies of different type II restriction enzymes in the presence high positive supercoiling and no supercoiling. The data for EcoRV is reproduced from Figure 3C. Error bars represent standard deviation (σ).

As stated previously, cleavage assays were not carried out in the optimal reaction buffers recommended by New England Biolabs. Therefore, only the relative change in cleavage efficiency between supercoiled states was considered for each enzyme.

To ensure that these cleavage efficiencies were not influenced by laser or flow induced DNA breakage, we repeated these experiments in the absence of any restriction enzymes. In the presence of a high degree of positive supercoiling, we observed that 4.8 ± 2.1% (n = 8) of molecules were cleaved/broken. In the absence of supercoiling, we observed 3.9 ± 1.3% (n = 8) of molecules were cleaved/broken. These results indicate that flow and laser related breakage does not contribute to overall enzyme-induced cleavage efficiency values significantly.

## Discussion

Our findings reveal a relationship between the catalytic activity of DNA-bending restriction enzymes and supercoiling. While our finding that the activity of EcoRV can be modulated by the topology of DNA is in line with previously published work, the use of single-molecule fluorescence microscopy allows us to expand upon this. Using fluorescently labelled EcoRV has allowed us to separate the binding activity of EcoRV from the cleavage activity of EcoRV. We have determined that DNA topology does not influence the binding of EcoRV, only the cleavage activity. Our observation is in line with the known mechanism of EcoRV binding, as it has been reported that EcoRV binds all DNA sequences with equal affinity however, cognate sequences are bent and cleaved upwards of a million times more readily than non-cognate sequences ^8,11,45^.

The increased cleavage efficiency of EcoRV that we observe in the presence of supercoiling aligns with previous work ^20^. However, the previous study proposes that EcoRV and several other restriction enzymes were able to locate their target sites through both 1D and 3D searching pathways. The 3D pathways were suggested to be assisted by plectonemes bringing distant sites in closer proximity ^20^. Interestingly, ClaI and PstI, restriction enzymes that are known to not induce a DNA-bend upon binding, were amongst the enzymes found to have an increased cleavage efficiency in the presence of supercoiled DNA. This is in contrast to what we observe for HindIII. A crucial difference between our method and the previous work, is that our DNA is stretched out, while the DNA in the previous study is not. Therefore, the effect of plectoneme-assisted 3D searching pathways would be reduced in our assay. The fact that EcoRV and KpnI, enzymes that bend DNA, still show enhanced activity under our conditions suggests that supercoiling facilitates enzyme activity through another mechanism, as well as 3D target searching.

As both EcoRV and KpnI are known to induce a significant DNA bend upon binding we hypothesise that the presence of a plectoneme at the cognate sequence will lower the energy required for the restriction enzyme to bend and ultimately cleave the DNA. This would make the cleavage reaction more energetically favourable in the presence of supercoiling. EcoRV bends the DNA to a greater extent than KpnI ^7,44^, potentially explaining the difference in cleavage efficiency under the same supercoiling conditions (Figure 4C). As HindIII only induces a minimal bend upon binding to DNA ^43^, the effect of supercoiling on its activity would be minimal. Therefore, this could explain why we do not see any supercoiling-dependent differences in cleavage efficiency for HindIII.

Our hypothesis that the presence of supercoiling may provide an energetic advantage to restriction enzymes that induce a significant DNA bend prior to cleavage is further supported by tension-dependent DNA cleavage studies ^46,47^. Using optical tweezers, differing forces were applied to DNA in order to determine the effect that tension has on the cleavage efficiency of both DNA-bending and non-DNA-bending restriction enzymes. It was found that, at forces of 20 and 40 pN, clear inhibition of EcoRV was observed. At these same forces, enzymes that do not bend DNA significantly, BamHI and EcoRI ^48,49^, were only very mildly inhibited. It was hypothesised that the application of tension to a DNA substrate lowers the cleavage efficiency of DNA-bending enzymes due to the increase in free energy change required for the enzyme to bind to and bend the DNA. Conversely, the presence of a plectoneme at the cognate sequence of a DNA-bending enzyme would lower the free energy change required for binding and bending to occur.

In our single-molecule supercoiling assay, the intercalating dye, SYTOX orange, was used to induce supercoiling upon topologically constrained DNA templates. We observed that the binding of SYTOX orange did not only induce supercoiling, but supercoiling then modulated the binding of SYTOX orange dye molecules (Figure 1D). We hypothesise that this binding modulation by supercoiling may not be limited to intercalating dyes such as SYTOX orange and, instead, supercoiling may be modulating other processes that involve intercalation between DNA base pairs. A number of proteins interact with DNA through intercalation, a specific example is SOX-4, a transcription factor that promotes tumorigenesis and tumour progression ^50–52^. Additionally, it has been reported that amino-acid side chains can intercalate between base pairs to distort the DNA helix, a process that is important for multi-protein complex assembly, protein recruitment, and transcription regulation ^53,54^. The modulation of intercalation by supercoiling that we have observed in our assay suggests that supercoiling may also be able to regulate the intercalation of these transcription factors and other proteins. While it is known that supercoiling is present in all processes that act on DNA, these findings suggest that DNA itself may be playing a more active role in modulating the processes acting upon it than what is currently thought.

While we have observed supercoiling modulating restriction enzyme activity by lowering the energy required for bending, it is likely that supercoiling similarly modulates other processes that act upon DNA, such as origin recognition ^55^, transcription ^56,57^, nucleosome assembly ^58^, and DNA repair ^59,60^. This presents the need for further research to determine the extent to which DNA itself modulates the processes acting upon it.

## Acknowledgements

We thank Jacob Lewis, Nicholas Dixon, Cees Dekker, Jaco van der Torre, Victoria Hill, Mahmoud Abdelhamid, and Alice Rhind-Tutt for helpful discussions and assistance in supercoiling assay development. We would also like to thank Aidan Fitch for developing and providing the PDMS block required for the cross-flow assay.

This work was supported by the Australian Research Council (DP210100067 to A.M.v.O.), the Australian National Health and Medical Research Council (Investigator grant 2007778 to L.M.S. and Investigator grant 1197069 to A.M.v.O.), The National Institutes of Health (RM1 GM130450 to A.M.v.O and RM1 GM13045 to L.M.S.), The Bruce Warren Molecular Horizons Early Career Fellowship (to S.H.M.), and an Australian Government Research Training Program Scholarship (to E.M.W.).

## Author contributions

Conceptualisation, E.M.W., L.M.S., A.M.v.O., T.D.C., and S.H.M.; Analysis, E.M.W., and S.H.M.; Funding Acquisition, L.M.S., and A.M.v.O.; Investigation, E.M.W., and S.H.M.; Methodology, E.M.W., L.M.S., A.M.v.O., and S.H.M.; Software, S.H.M., and L.M.S., Supervision, S.H.M., A.M.v.O, T.D.C., and L.M.S.; Validation, E.M.W.; Visualisation, E.M.W, L.M.S., A.M.v.O., and S.H.M.; Writing – Original Draft, E.M.W., and L.M.S.; Writing – Review and Editing, E.M.W., L.M.S., A.M.v.O., T.D.C., and S.H.M.

## Data availability

All data available at 10.5281/zenodo.15186176 and home-built ImageJ plugins on https://github.com/Single-molecule-Biophysics-UOW.

## Additional information

### Competing interests

The authors declare no competing interests

## Supplementary Figures

**Supplementary Table S1:**
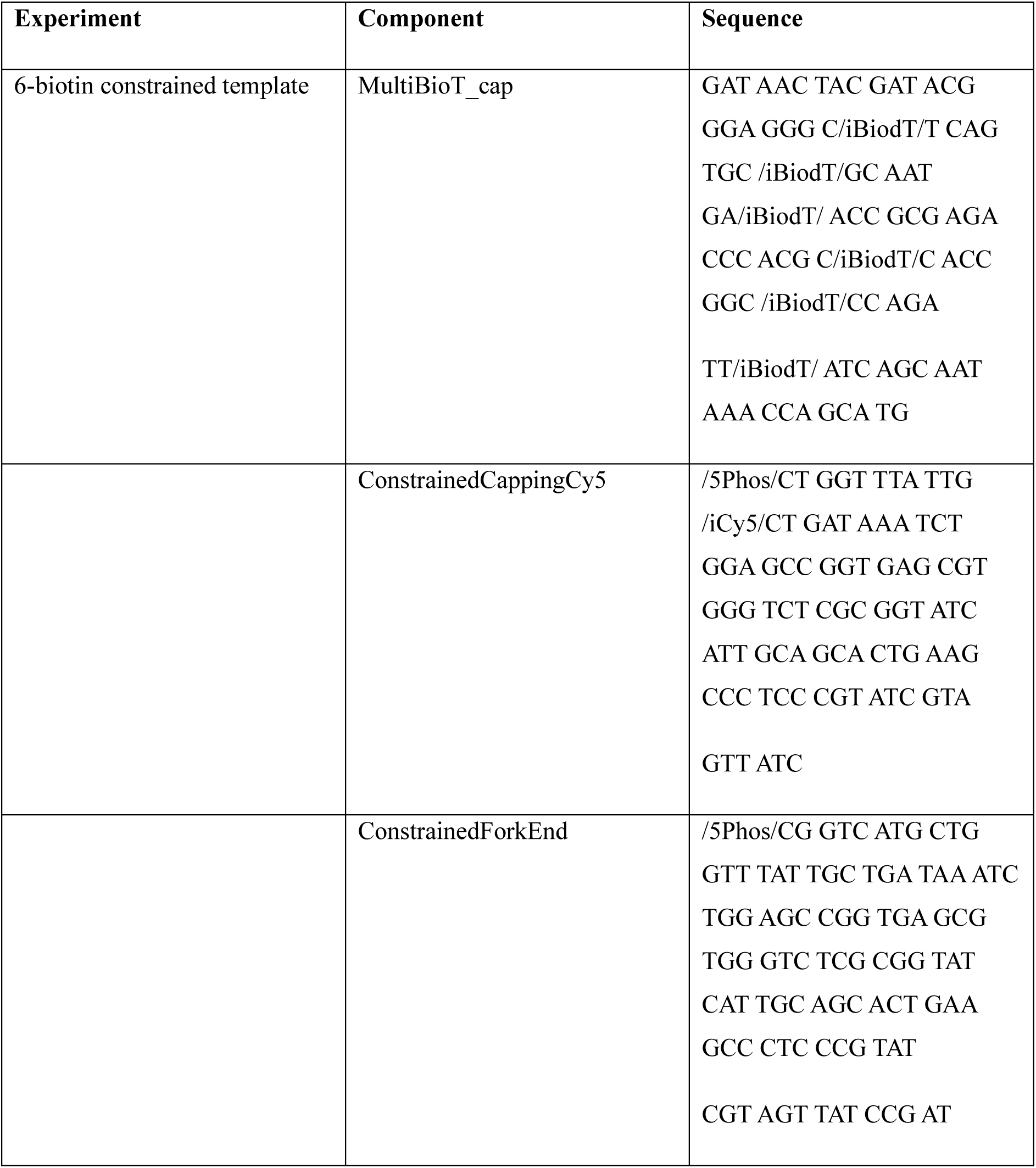
Oligonucleotide sequences used in this study. MultiBioT_cap was annealed to ConstainedCappingCy5 and ConstrainedForkEnd separately to construct two different oligo blocks.

**Supplementary Table S2:**
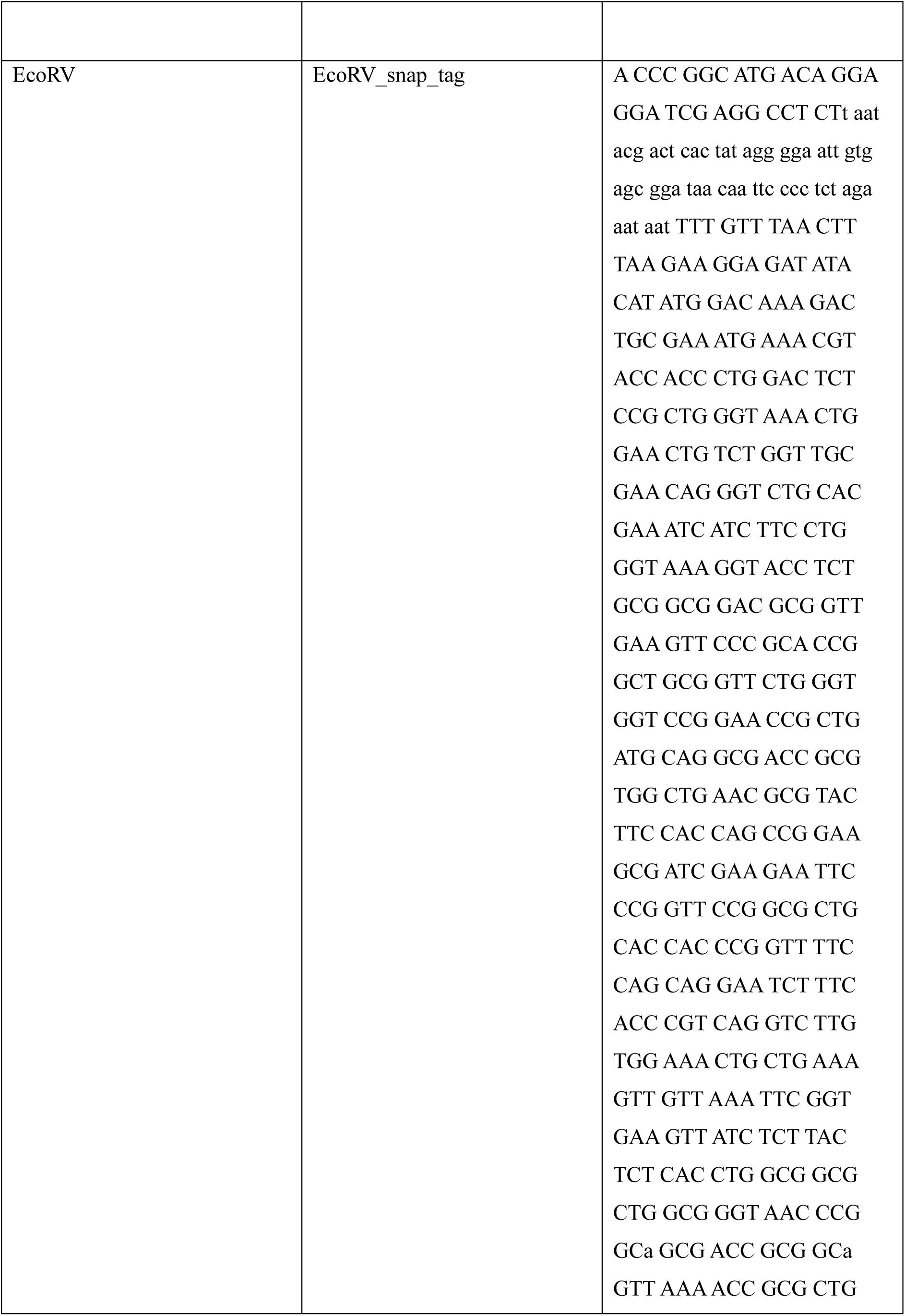
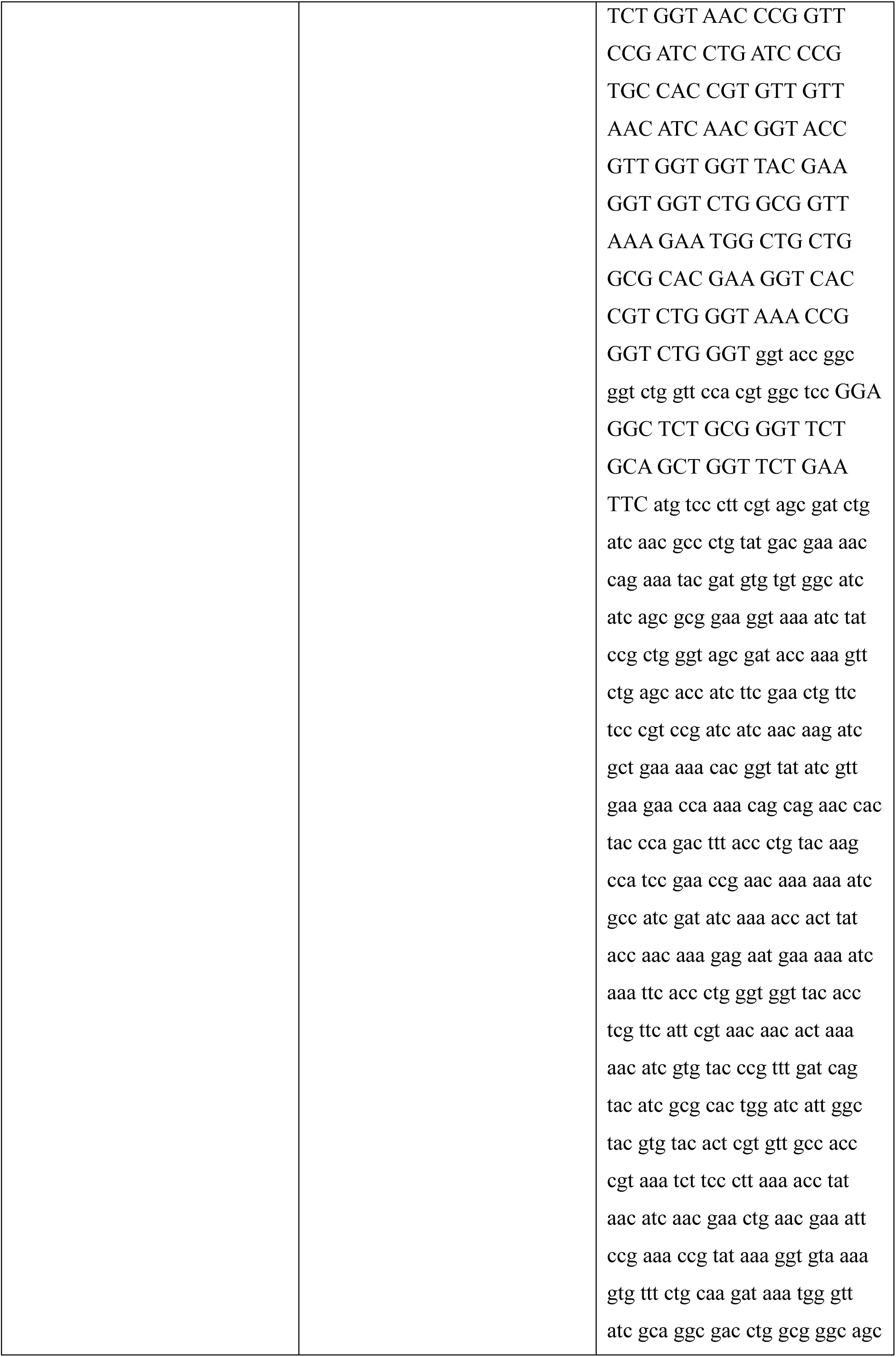
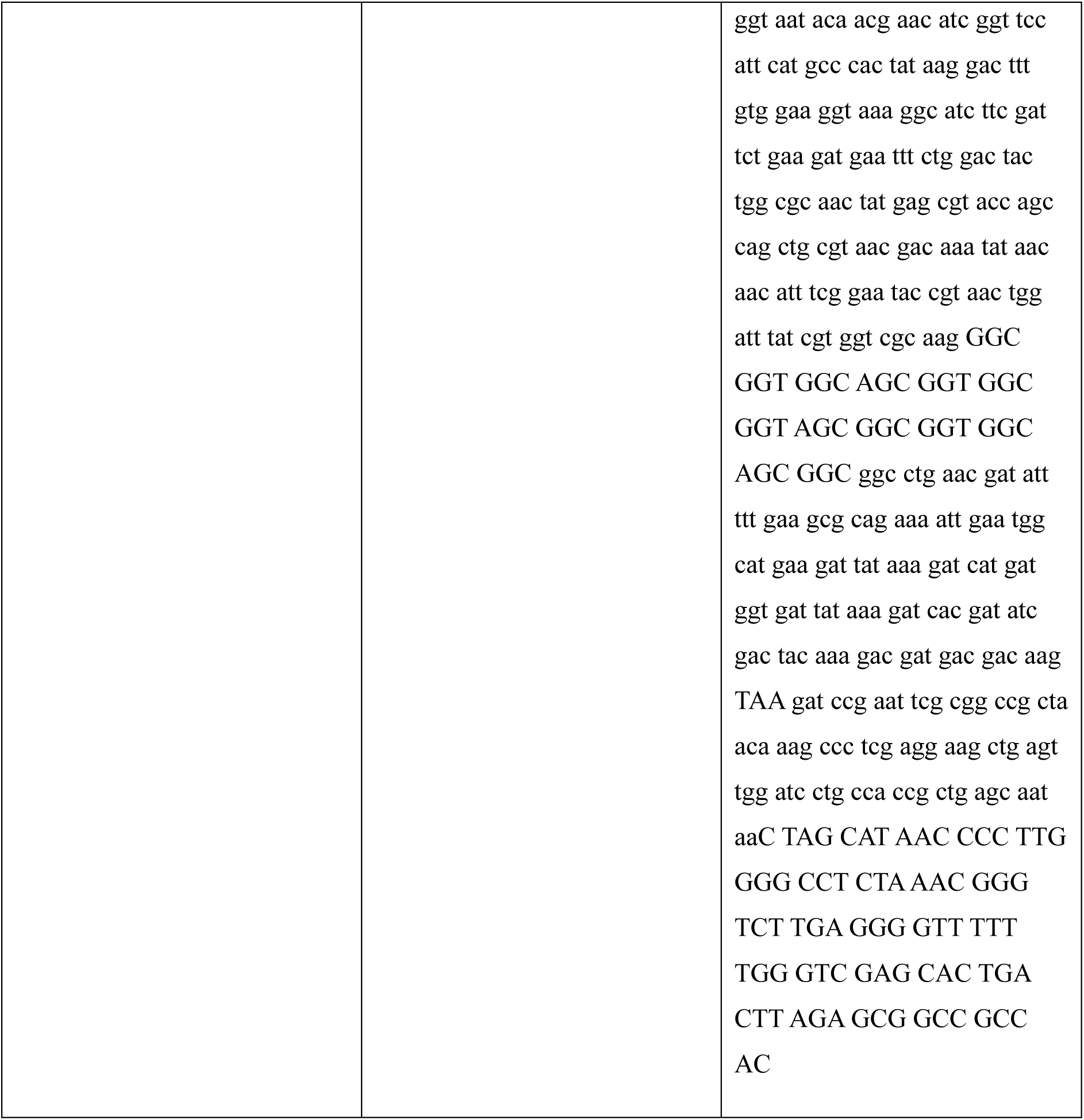
EcoRV sequence.

**Supplementary Figure S1:**
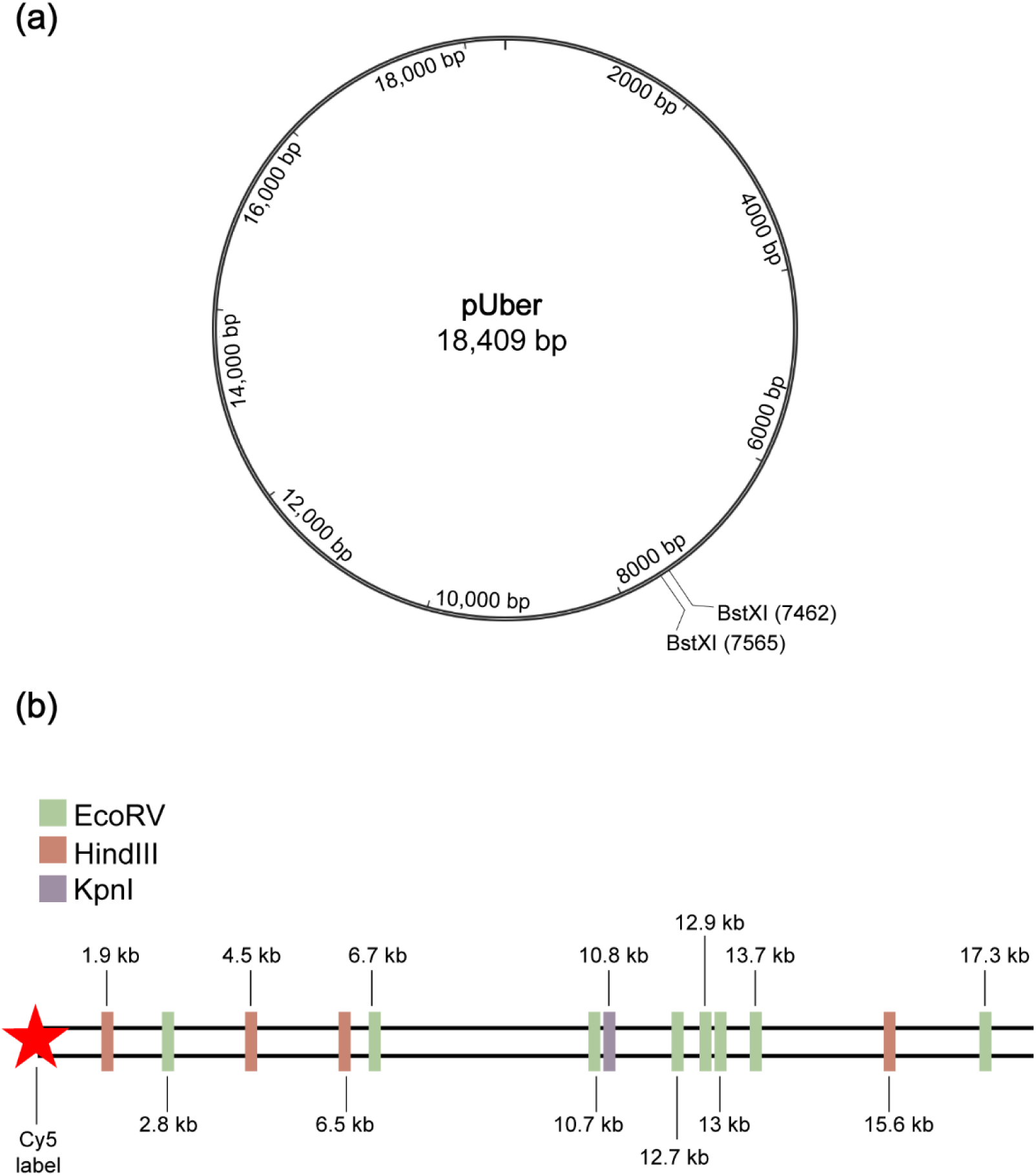
Plasmid map of pUber and location of EcoRV, HindIII, and KpnI restriction sites. **(A)** Plasmid map of the 18-kb plasmid pUber, highlighting the positions of the BstXI restriction sites used to linearize the plasmid for template DNA template construction. **(B)** Simplified schematic of the constrained DNA template produced from linearized pUber with the positions of the Cy5 end label and restriction sites for EcoRV, HindIII, and KpnI indicated.

**Supplementary Figure S2:**
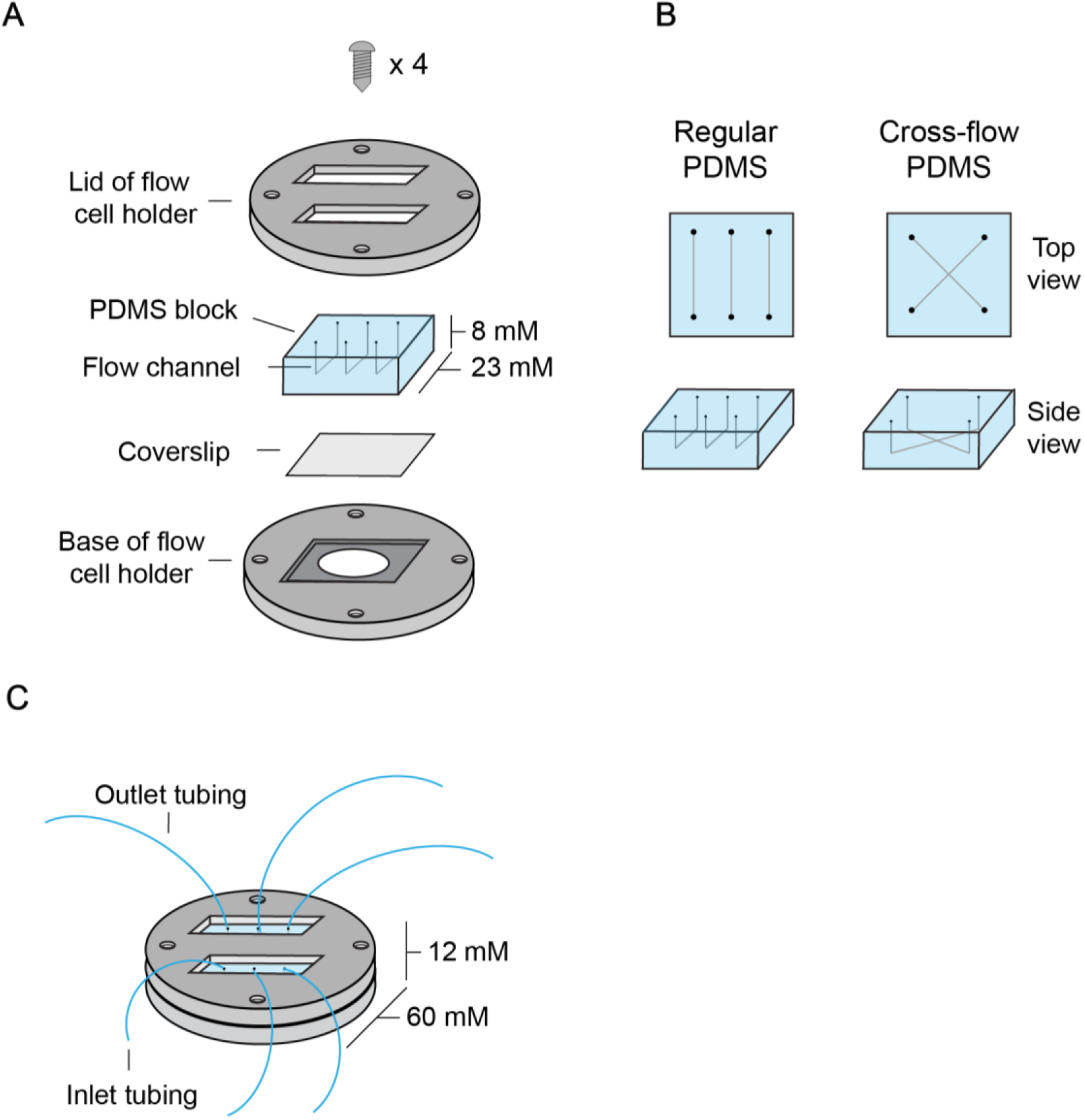
Flow cell construction. **(A)** Schematic of flow cell components and assembly. The PEG-biotin-functionalized microscope coverslip sits on top of the flow cell holder base. A PDMS block, either three channel or cross-flow, is placed on top of the coverslip. The flow cell holder lid is then secured to the top with 4 screws. **(B)** Schematics of the regular three channel PDMS block and the cross-flow PDMS block, highlighting the channels from top and side views. **(C)** Schematic of constructed flow cell with inlet and outlet tubing inserted.

**Supplementary Figure S3:**
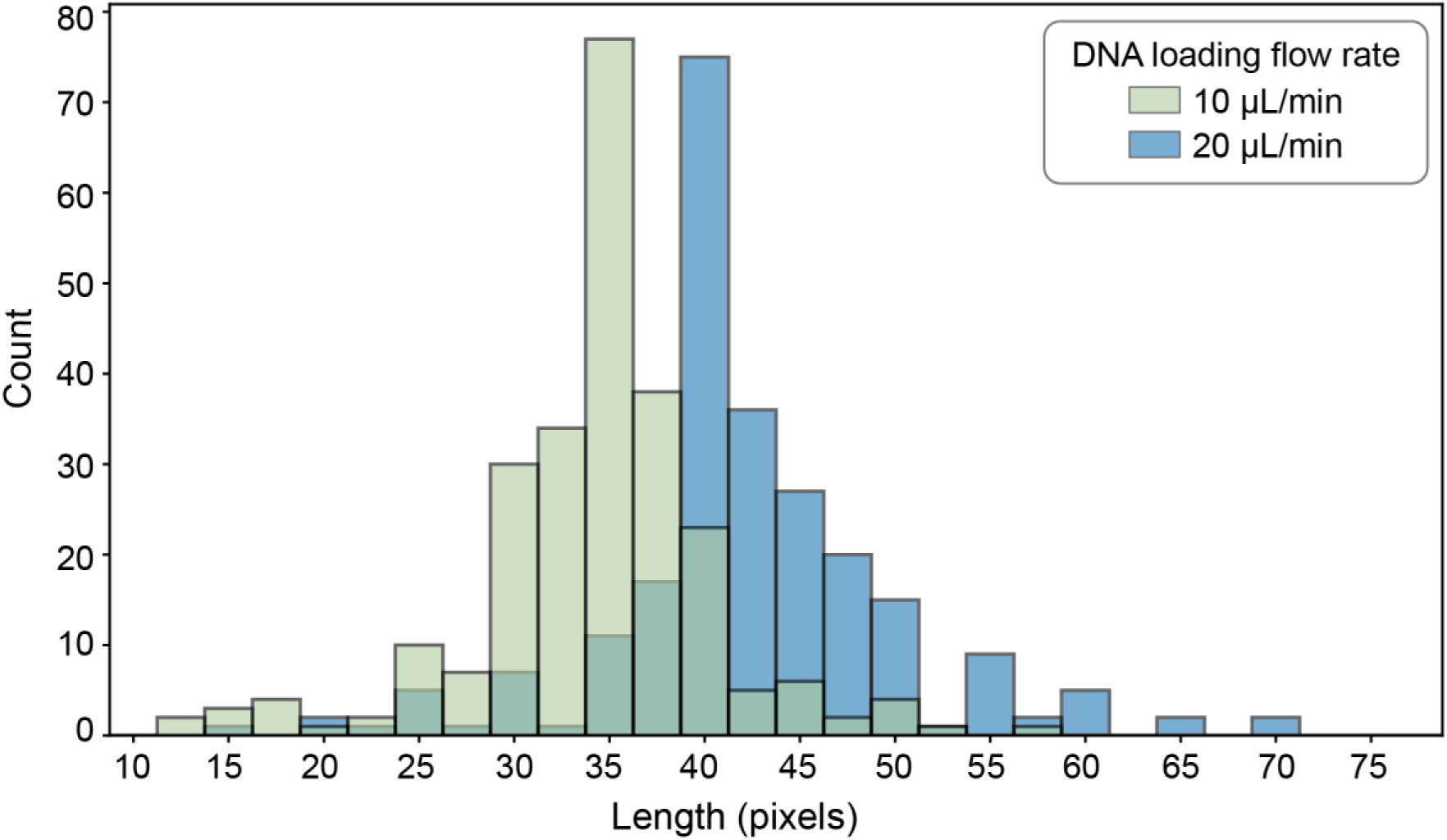
DNA stretch Analysis. Histogram showing distribution of DNA lengths in pixels for two different DNA-loading flow rates, 10 µL/min and 20 µL/min. When DNA is loaded into the flow cell at a flow rate of 10 µL/min, the average length of the DNA templates are 34.4 ± 6.3 pixels (n = 241). When DNA is loaded at a flow rate of 20 µL/min, the average length of the DNA templates are 42.7 ± 8.5 pixels (n = 250).

**Supplementary Figure S4:**
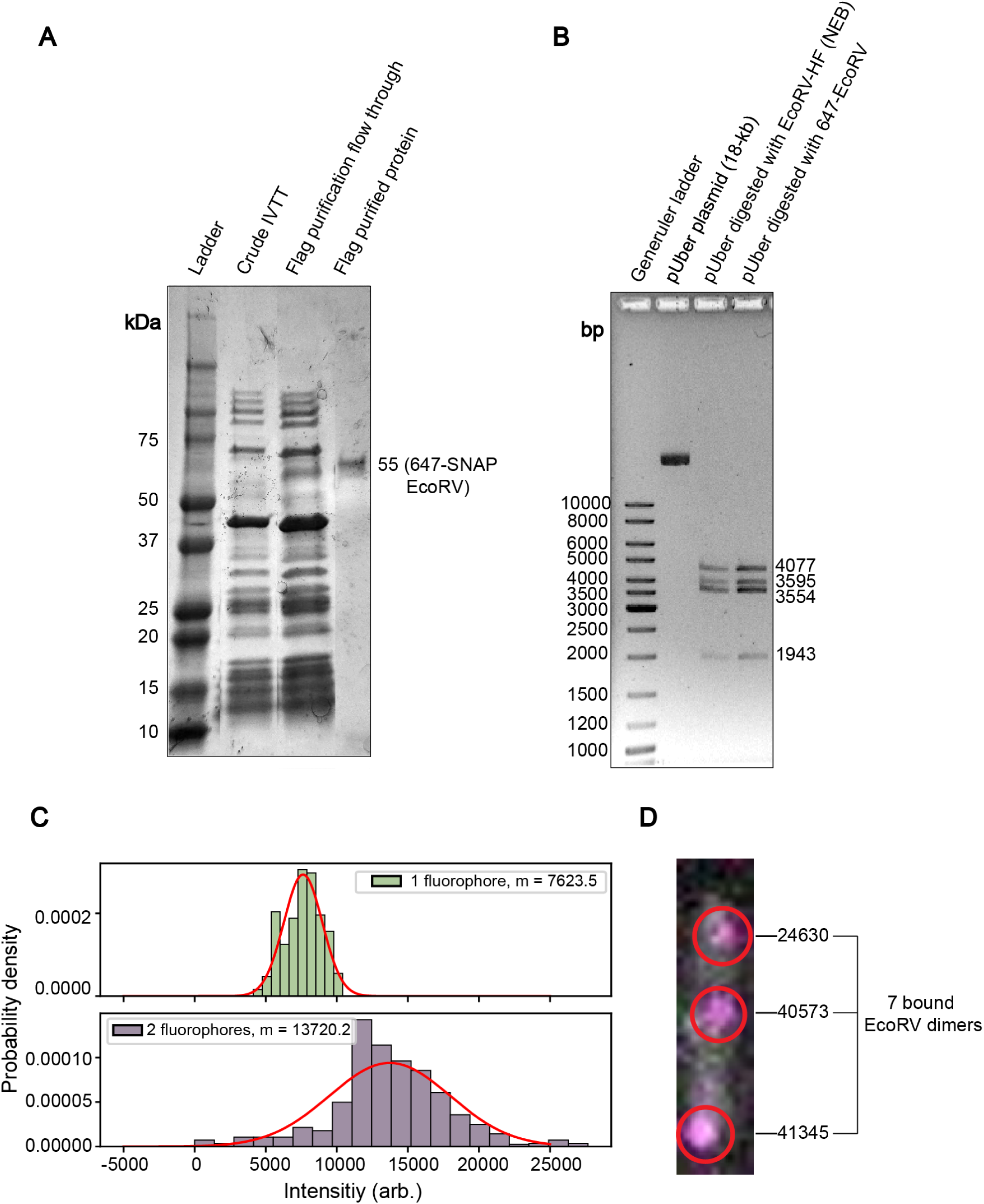
Cell-free expression and labeling of EcoRV. **(A)** Purification of EcoRV following cell-free expression with IVTT and fluorescent labeling. A 4–20% SDS-PAGE gel showing the sample after IVTT prior to flag purification (lane 2), the flag purification flow through (lane 3), and the sample following flag purification where a single band is present at the expected size of 55 kDa, indicating successful purification (lane 4). **(B)** 647-EcoRV activity test. 1% agarose gel showing the digestion of the 18-kb plasmid pUber with commercial EcoRV-HF (NEB) (lane 3) and 647-EcoRV (lane 4) following labeling and purification. Results indicate that 647-EcoRV is active and efficient. **(C)** Distribution of fluorescence intensities obtained through a photobleaching assay. The average intensity of one fluorophore (EcoRV monomer) is equal to 7623.5, and the average intensity of two fluorophores (EcoRV dimer) is equal to 13720.2. **(D)** Determining the number of EcoRV restriction sites occupied on a single DNA template. Average z-projections of each molecule were generated and the 647-nm signal that was colocalized with a DNA template was manually picked. The integrated intensity of each colocalized 647-nm spot was determined and the intensity for all spots on a single molecule were summed. This value was then divided by the average intensity of two 647 fluorophores, which corresponds to the intensity of an EcoRV dimer. From this, the number of EcoRV restriction sites occupied per molecule was determined.

**Supplementary Figure S5:**
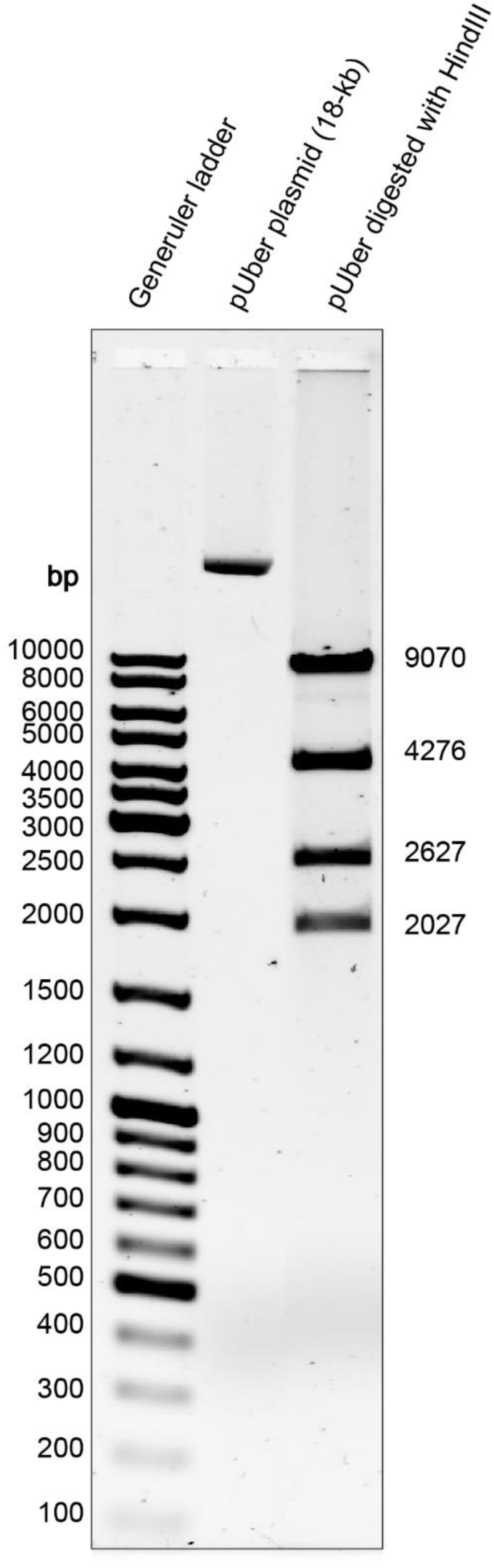
Gel-based HindIII activity test. An 18-kb circular plasmid (pUber) was digested with HindIII-HF (NEB) as per company instructions. Digested products were loaded on a 1% agarose gel to determine the cleavage efficiency of HindIII in an ensemble-based assay.

## Notes

### Competing Interest Statement

The authors have declared no competing interest.

